# Long-read assembly and comparative evidence-based reanalysis of *Cryptosporidium* genome sequences reveal new biological insights

**DOI:** 10.1101/2021.01.29.428682

**Authors:** Rodrigo P. Baptista, Yiran Li, Adam Sateriale, Mandy J. Sanders, Karen L. Brooks, Alan Tracey, Brendan R. E. Ansell, Aaron R. Jex, Garrett W. Cooper, Ethan D. Smith, Rui Xiao, Jennifer E. Dumaine, Matthew Berriman, Boris Striepen, James A. Cotton, Jessica C. Kissinger

## Abstract

Cryptosporidiosis is a leading cause of waterborne diarrheal disease globally and an important contributor to mortality in infants and the immunosuppressed. Despite its importance, the *Cryptosporidium* community still relies on a fragmented reference genome sequence from 2004. Incomplete reference sequences hamper experimental design and interpretation. We have generated a new *C. parvum* IOWA genome assembly supported by PacBio and Oxford Nanopore long-read technologies and a new comparative and consistent genome annotation for three closely related species *C. parvum*, *C. hominis* and *C. tyzzeri*. The new *C. parvum* IOWA reference genome assembly is larger, gap free and lacks ambiguous bases. This chromosomal assembly recovers 13 of 16 possible telomeres and raises a new hypothesis for the remaining telomeres and associated subtelomeric regions. Comparative annotation revealed that most “missing” orthologs are found suggesting that species differences result primarily from structural rearrangements, gene copy number variation and SNVs in *C. parvum, C. hominis* and *C. tyzzeri*. We made >1,500 *C. parvu*m annotation updates based on experimental evidence. They included new transporters, ncRNAs, introns and altered gene structures. The new assembly and annotation revealed a complete DNA methylase *Dnmt2* ortholog. 190 genes under positive selection including many new candidates were identified using the new assembly and annotation as reference. Finally, possible subtelomeric amplification and variation events in *C. parvum* are detected that reveal a new level of genome plasticity that will both inform and impact future research.

## INTRODUCTION

*Cryptosporidium spp.* are parasitic apicomplexans that cause moderate-to-severe diarrhea in humans and animals. Studies funded by the Bill and Melinda Gates Foundation, revealed that *Cryptosporidium* is one of the most common causes of waterborne disease in humans and the second leading cause of diarrheal etiology in children < 2 years resulting in ~60,000 fatalities worldwide (Kotloff et al. 2013; Collaborators 2017). In 2016, acute infections caused more than 48,000 global deaths and more than 4.2 million disability-adjusted life years lost (Khalil et al. 2018).

Currently, 38 species of *Cryptosporidium* are recognized by the scientific community (Slapeta 2013; Feng et al. 2018). Most are host-adapted, and host species range from fish to mammals. Of these, 15 species have had their genome sequence generated and assembled however, only 8 are annotated. Most genomic sequence data are from the zoonotic *C. parvum* and anthroponotic *C. hominis,* the species primarily detected in humans (Chalmers et al. 2011; Zahedi et al. 2016; Khan et al. 2017). These two species are only 3-5% divergent at the DNA level (Mazurie et al. 2013).

As the *Cryptosporidium* field is exploding with new-found interest and much needed breakthroughs in genetics and culturing (Vinayak et al. 2015; Morada et al. 2016; DeCicco RePass et al. 2017; Heo et al. 2018; Wilke et al. 2019), the limitations of existing reference genome sequences need to be addressed. The *C. parvum* IOWA II reference genome sequence was assembled with limited physical map data (Abrahamsen et al. 2004) and experimental data for training gene finders and providing functional annotation were limited to a few hundred ESTs from oocysts and sporozoite stages only. Genomic, transcriptomic and proteomic work on this important pathogen has been lacking due to the obligate quasi-intracellular nature of portions of the parasite’s life cycle, the historical lack of a continuous *in vitro* tissue culture system, the parasite’s small size relative to host cells and difficult animal models. The physical map for the *C. parvum* IOWA II reference assembly was generated from two different studies that utilized the genome-wide HAPPilly anchored physical mapping technique, an *in vitro* linkage technique based on screening approximately haploid amounts of DNA by PCR, which is very accurate (Piper et al. 1998; Bankier et al. 2003). Even with these cutting-edge approaches at the time, some regions, especially chromosome ends, lacked support or were poorly resolved. Subsequent whole genome sequencing data often remain unassembled or in a large number of contigs.

In 2015, the reference genome sequence of *C. parvum* was re-annotated based on new RNA-seq evidence and a new *C. hominis* sequence from a recent human isolate (UdeA01) was generated (Isaza et al. 2015). Many ambiguities in gene models were improved based on the new RNA-seq data, but since the new *C. hominis* UdeA01 genome is still fragmented and the annotation was primarily based on the 2004 *C. parvum* IOWA II reference annotation. Additionally, annotation of sequences from closely related species has been performed independently and are not consistent, causing possible misinterpretations regarding gene content and species-specific genes.

Incomplete and misassembled (i.e. gapped sequence, indels, frameshifts, compressed repetitive regions, inversions) reference genome sequences such as those shown in (Guo et al. 2015) can mislead interpretations of the differences between isolates and species resulting in extra assays to confirm insertions, deletions and copy number variations (CNVs). Since incomplete and misassembled sequences are usually caused by repetitive and complex sequence regions, it is imperative to revisit older reference genome sequences with new long-read technologies to close gaps and expand regions of the genome sequence that were misassembled or collapsed into shorter regions because they are repetitive. Long-read sequence technologies (PacBio and Oxford Nanopore) are becoming an essential tool to close full genome sequence assemblies across the tree of life (Vembar et al. 2016; Diaz-Viraque et al. 2019; Miga et al. 2020). They can be used to resolve complex regions such as repetitive content, structural variants (SVs) such as inversions, translocations and duplications, or for use as scaffolding evidence for existing fragmented genome assemblies (Mahmoud et al. 2019). They are proving crucial for completing assemblies of pathogen genome sequences that are often riddled with large virulence-related gene families that have been collapsed or improperly assembled in shorter-read assemblies. Here we provide a new *de novo* reference long-read assembly for *C. parvum* strain IOWA (DNA obtained from the ATCC) and new consistent, comparative genome annotations for *C. parvum* IOWA-ATCC, *C. hominis* UdeA01 and *C. tyzzeri* UGA55.

## RESULTS

### An improved long-read based genome assembly for *Cryptosporidium parvum* (IOWA-ATCC)

The current *C. parvum* IOWA II reference genome assembly, generated in 2004, is good, but it still has 10 gapped regions of unknown size, 14,600 ambiguous bases, and is missing 6 telomeres. By aligning Illumina reads against this reference sequence, we have detected many collapsed regions, suggesting misassembled repetitive and complex regions (Supplemental Table S1). To resolve these issues, we generated a new PacBio+Illumina+Nanopore hybrid genome assembly for the *Cryptosporidium parvum* strain IOWA (ATCC^®^PRA-67DQ™) with DNA from oocysts/sporozoites purchased from ATCC. To minimize strain variation differences, we performed our analysis on the same strain, however because there is a 14-year time window of propagation between these two isolates, and cryopreservation has only been recently made possible (Jaskiewicz et al. 2018), we modified the strain name to IOWA-ATCC.

The new *C. parvum* IOWA-ATCC genome statistics are compared to the current *C. parvum* IOWA II reference genome sequence and *C. hominis* 30976 and *C. tyzzeri* UGA55 two closely related species with different host preferences and pathogenicity (Slapeta 2013; Nader et al. 2019; Sateriale et al. 2019) (Table 1). These particular *C. hominis* and *C. tyzzeri* assemblies were selected because they are the best available. The new long-read assembly increases the genome size by 19,939 bases and identifies 13 of 16 expected telomeres. There are no gaps and no ambiguous bases. As expected, the *C. parvum* IOWA-ATCC genome sequence has diverged slightly but shares 99.93% average pairwise identity with the 2004 assembly in regions that exist in both assemblies (Supplemental Table S2). The main *Cryptosporidium* subtyping marker, the 60 kDa surface protein (*gp60* locus subtype IIa) shows 4 amino acid differences between the IOWA-ATCC and 2004 assemblies (Supplemental Fig. S1).

**Table 1.**
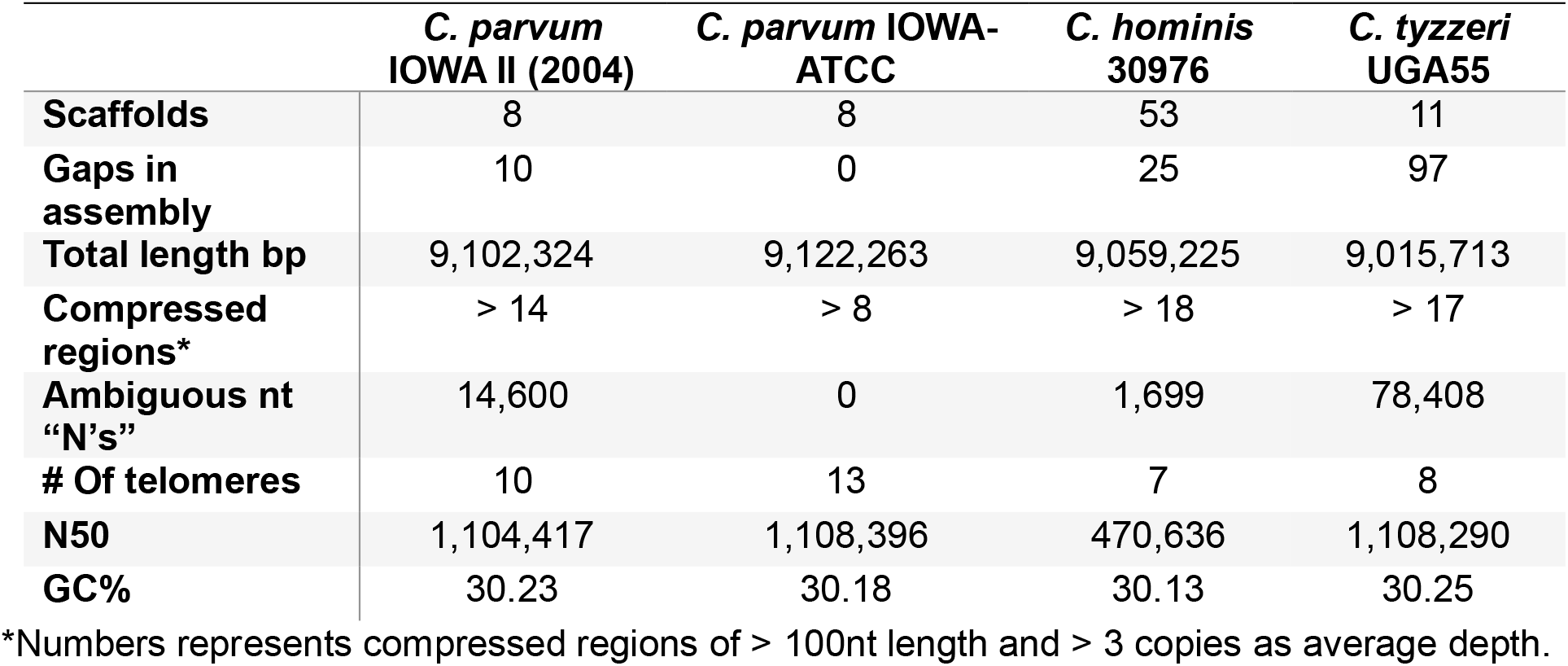
Cryptosporidium Genome Assembly Statistics.

|  | *C. parvum*<br>IOWA II (2004) | *C. parvum* IOWA-ATCC | *C. hominis*<br>30976 | *C. tyzzeri*<br>UGA55 |
| --- | --- | --- | --- | --- |
| **Scaffolds** | 8 | 8 | 53 | 11 |
| **Gaps in**<br>**assembly** | 10 | 0 | 25 | 97 |
| **Total length bp** | 9,102,324 | 9,122,263 | 9,059,225 | 9,015,713 |
| **Compressed**<br>**regions\*** | > 14 | > 8 | > 18 | > 17 |
| **Ambiguous nt**<br>**"N's"** | 14,600 | 0 | 1,699 | 78,408 |
| **# Of telomeres** | 10 | 13 | 7 | 8 |
| **N50** | 1,104,417 | 1,108,396 | 470,636 | 1,108,290 |
| **GC%** | 30.23 | 30.18 | 30.13 | 30.25 |

\*Numbers represents compressed regions of > 100nt length and > 3 copies as average depth.

### Structural differences between the *C. parvum* IOWA assemblies

The 2004 *C. parvum* IOWA II genome assembly used Sanger reads combined with available HAPPY-map data to scaffold the contigs. We compared the 2004 and IOWA-ATCC assemblies to identify potential rearrangements. Small and large rearrangements were detected primarily in chromosomes 2, 4 and 5 (Fig. 1A). Chromosomal inversions may be assembly artifacts or represent genuine differences generated during evolution. Inversions are often associated with speciation events (de Meeus et al. 1998; Rieseberg 2001; Nosil and Feder 2012). We thus investigated the synteny between *C. parvum* IOWA II and ATCC, *C. hominis* 30976 and *C. tyzzeri* UGA55 and observed that *C. hominis* and *C. tyzzeri* also share the large inversions in their chr 4 and chr 5. Examination of the inverted region boundaries revealed that sequences in these regions in the 2004 *C. parvum* assembly consist of ambiguous nucleotide bases or physical gaps (Fig. 1B). These results suggest that the 2004 *C. parvum* assembly may contain misassembled scaffolds, but the data do not rule out their presence in that isolate. Better assemblies will be needed for the other isolates to determine the true level of synteny across these species.

**Figure 1.**
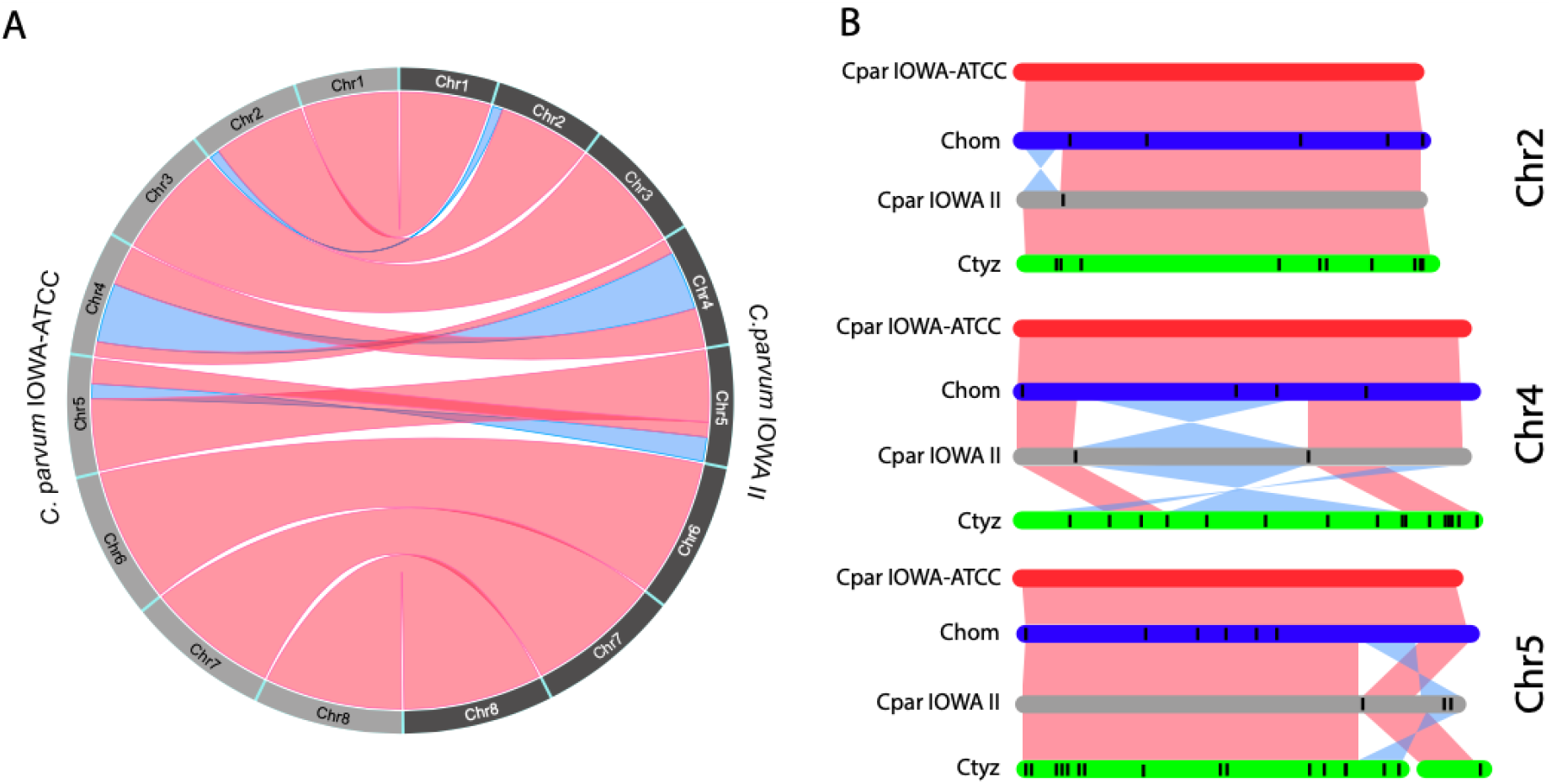
Syntenic relationships and inversions detected between the *Cryptosporidium* genome assemblies. (A) Circos plot of synteny between *C. parvum* IOWA-ATCC and IOWA II. (B) Synteny between chromosomes 2, 4 and 5 of *C. hominis* 30976, *C. parvum* IOWA II and *C. tyzzeri* UGA55. Each vertical black line within a chromosome represents a known gap region. Syntenic regions between chromosomes are shown in red and inverted regions are represented in blue. Cpar: *C. parvum*; Chom: *C. hominis*; Ctyz: *C. tyzzeri*.

### New consistent annotation across*Cryptosporidium* species provides insights

We consistently annotated and compared the three closely related, yet biologically different, *Cryptosporidium* species (genome identity > 95%) to assess differences in gene content. The new annotation for each species was generated with three *de novo* approaches and evidence-based manual annotation. Curation of the annotation was performed in a 3-way comparison between each pair of genome sequences to take full advantage of syntenic regions. The comparison permitted the use of data from one species to assess computational predictions in the others. By following this approach, fragments of genes that were previously missed in *C. hominis* were identified, permitting a more accurate identification of genuinely shared and species-specific genes in these species. This approach resulted in > 1500 gene structure alterations leading to an improved functional annotation. The changes increased the number of predicted genes, introns and exons (Table 2). The average mRNA length increased due to complete coding sequences (CDS) and the addition of exons to form larger genes. Notably, these structural fixes led to the repair of several genes, including finding and correcting the N-terminus of the DNA methylase ortholog, *Dnmt2* (Supplemental Fig. S2).

**Table 2.** Reannotation Summary Statistics.

|  | *C. parvum* IOWA II |  |  | *C. hominis* |  | *C. tyzzeri* UGA55 |
| --- | --- | --- | --- | --- | --- | --- |
|  | IOWA II Before | IOWA II After | IOWA-ATCC | UdeA01* "Before" | 30976 "After" | New |
| Total sequence length (bp) | 9,102,324 | 9,102,324 | 9,122,263 | 9,043,938 | 9,059,225 | 9,015,884 |
| Number of genes | 3,886 | 4,020 | 3,954 | 3,863 | 3,996 | 4,037 |
| Number of CDS | 3,805 | 3,944 | 3,944 | 3,818 | 3,959 | 3,986 |
| Number of exons | 4,104 | 5,043 | 5,322 | 4,546 | 5,045 | 5,136 |
| Number of introns | 238 | 1,020 | 1,371 | 683 | 1,040 | 1,089 |
| Shortest intron (bp) | 9 | 36 | 36 | 36 | 36 | 22 |
| Pseudogenes | 74 | 114 | 1 | 45 | 88 | 62 |
| % of genome covered by CDS | 75.4 | 82.1 | 79.53 | 76.1 | 83.6 | 79.2 |

\*A previous annotation does not exist for the *C. hominis* 30976 sequence, so the results are compared to the recently annotated *C. hominins* UdeA01 strain.

*Cryptosporidium* has a very compact genome with < 20% being intergenic. As a result, RNA-Seq data, which is the best evidence for annotation, contains reads that overlap adjacent genes creating false fusions of exons belonging to different genes. Available strand-specific RNA-seq was used to characterize some of these regions but expression data were not available for all predicted genes, thus, genes of unknown function in close proximity on the same strand remain problematic. The expression data also revealed alternative splicing and potential non-coding RNAs (ncRNAs) predominantly anti-sense lncRNAs with differential expression (Li et al. 2020).

### Functional annotation

Several approaches to assess function were applied including InterPro scan and I-TASSER among others (see methods). 138 new protein annotations were generated or modified, the rest are unchanged. The percentage of *C. parvum* genes annotated as uncharacterized proteins was reduced from 40% to 33% in all reannotated sequences (Supplemental Table S3). Many new features including domain and repeat content were added to 738 previously uncharacterized proteins. 729 predicted *C. parvum* CDSs have signal peptides and 1990 have GO assignments. 1414 CDSs were further assessed for confidence using I-TASSER protein structure searches and 1008 predicted structures were assigned as high-confidence by random forest categorization (Supplemental Table S4). 143 previously uncharacterized proteins were assigned with high confidence GO terms. The top functional annotation terms observed following re-annotation were protein kinases, AAA+ATPases, TRAP, DEAD/DEAH box proteins, Ras GTPases, WD40-repeat containing proteins, ABC transporters, RNA recognition motifs, Palmitoyltransferases and insulinase-like proteases.

### New transporters were detected and annotated

Following functional annotation, we further characterized the newly identified transporter genes using three different prediction methods. A total of 145 proteins in *C. parvum* IOWA-ATCC and *C. hominis* 30976 were identified as transporters including 128 confident candidates and 24 putative candidates (Supplemental Table S5). This represents an increase of 53 transporters relative to the *C. parvum* IOWA II GO annotation (CryptoDB Release 36) and an increase of 93 relative to TransportDB v2.0 (http://www.membranetransport.org/transportDB2/index.html). The predicted transporters in *Cryptosporidium* are mostly related to purine metabolism, peptidoglycan biosynthesis, oxidative phosphorylation and N-Glycan biosynthesis pathways (Fig. 2). Six translocases were also identified.

**Figure 2.**
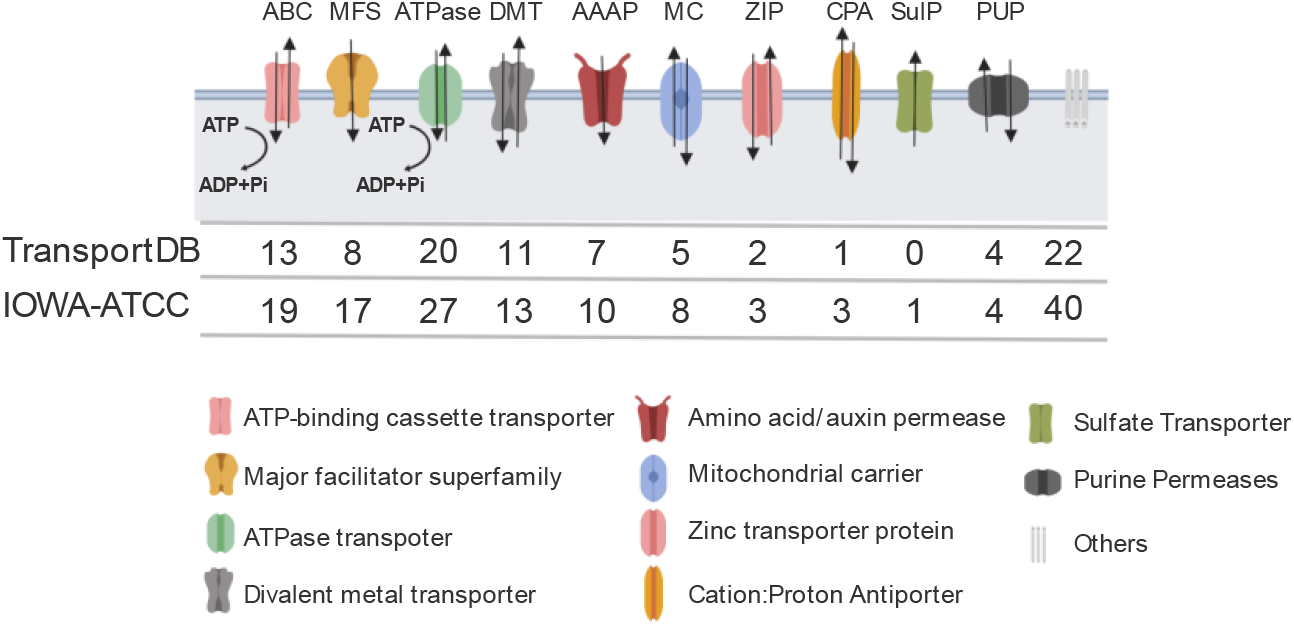
Reannotation reveals new transporters in *Cryptosporidium parvum* IOWA-ATCC. Numbers of transporters corresponds to the counts of genes encoding each type of transporter protein. ABC: ATP-binding cassette transporter; MFS: Major facilitator superfamily; DMT: Divalent metal transporter; AAAP: amino acid/auxin permease; MC: mitochondrial carrier; ZIP: Zinc transporter protein; CPA: Cation/Proton Antiporter; SulP: Sulfate Transporter; and PUP: Purine Permeases.

### Comparative analysis of closely related species of *Cryptosporidium*

*Cryptosporidium* species have a broad host spectrum with most species being largely host-adapted with a few zoonotic exceptions, principally *C. parvum*. Yet, despite these differences, there is a cluster of species with high synteny relative to other species outside of this cluster (Fig. 3; Supplemental Table S6).

**Figure 3.**
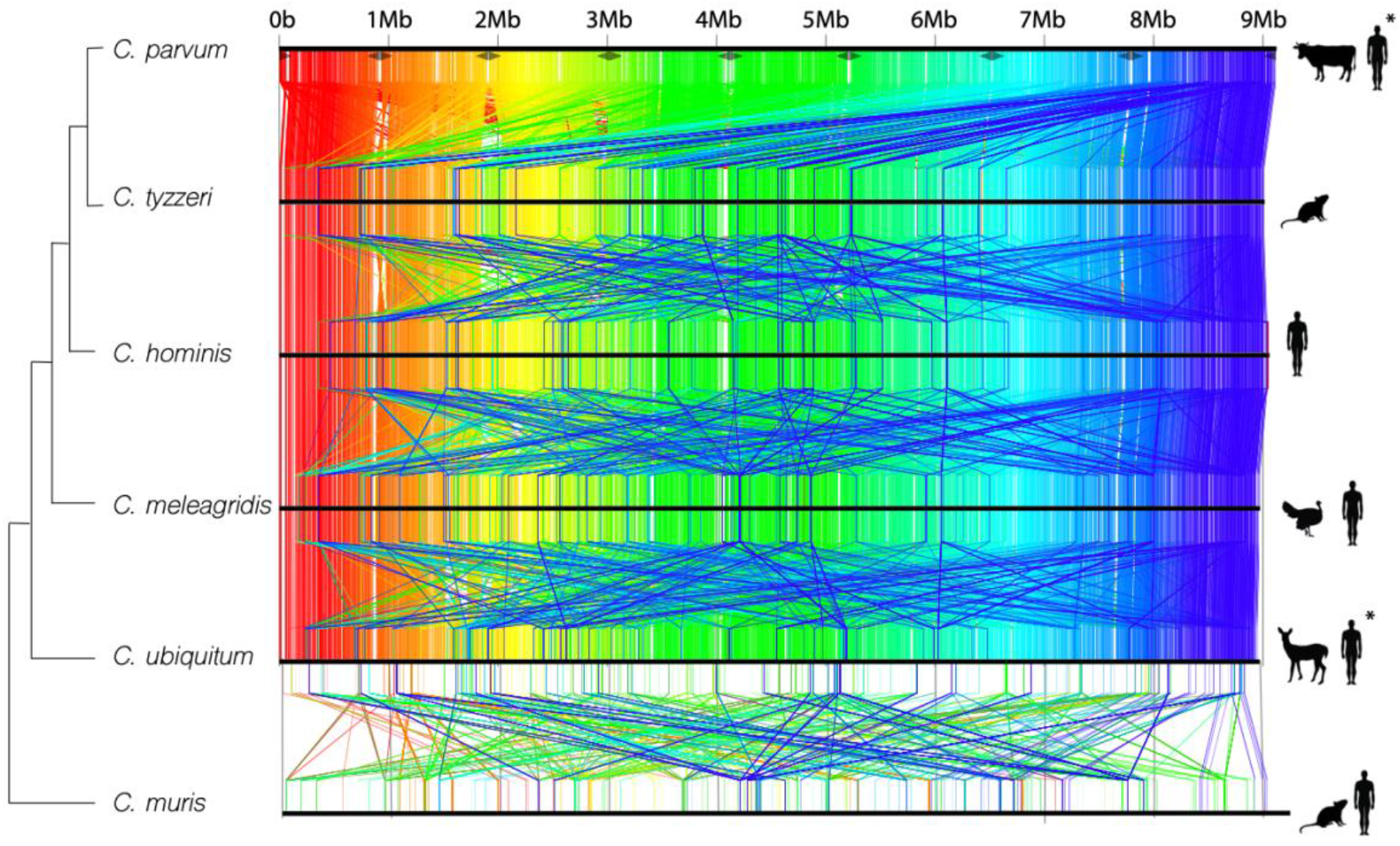
Comparative genome-wide synteny between six different species of *Cryptosporidiu*m. Highly conserved regions between the genomes are colored in order from red (5′ end of chromosome 1) to blue (3′ end of chromosome 8) with respect to genomic position of *C. parvum*. The cladogram topology was determined via a maximum likelihood analysis of 2700 revisited single copy orthologs. Animal icons represent the major hosts for these species. \**C. parvum* and *C. ubiquitum* are zoonotic with many hosts.

The consistent annotation of the species closest to *C. parvum* IOWA-ATCC, *C. hominis* 30976 and *C. tyzzeri* UGA55, permitted the detection of differences in protein encoding gene content and copy number variation. An automated orthology analysis between all three gene sets revealed that, ~94% of the genes were conserved among all species. Of the 4,008 ortholog groups identified, most annotated gene families were maintained with a similar number of paralogs (max = 6) detected in the same ortholog group, but the number of singletons varied between the three species (Fig. 4A; Supplemental Table S7). Some of these post-comparative annotation gene differences appeared to be unique to a particular species (Supplemental Table S8). Of the 224 singletons detected, we observed only 0, 1 and 1 potential truly species-specific genes in *C. parvum* IOWA-ATCC, *C. hominis* 30976 and *C. tyzzeri* UGA55, respectively following manual inspection (Fig. 4B). Both species-specific genes are uncharacterized proteins. The remaining 253 singletons are detected but incomplete in the fragmented assemblies of *C. hominis* and *C. tyzzeri*, appearing as split genes, frame-shifts, missed calls near a gap or contig break and putative false gene predictions in small contigs (Fig. 4C; Supplemental Fig. S3). The majority of gene content differences between these species are gene copy number variations and not gene presence or absence.

**Figure 4.**
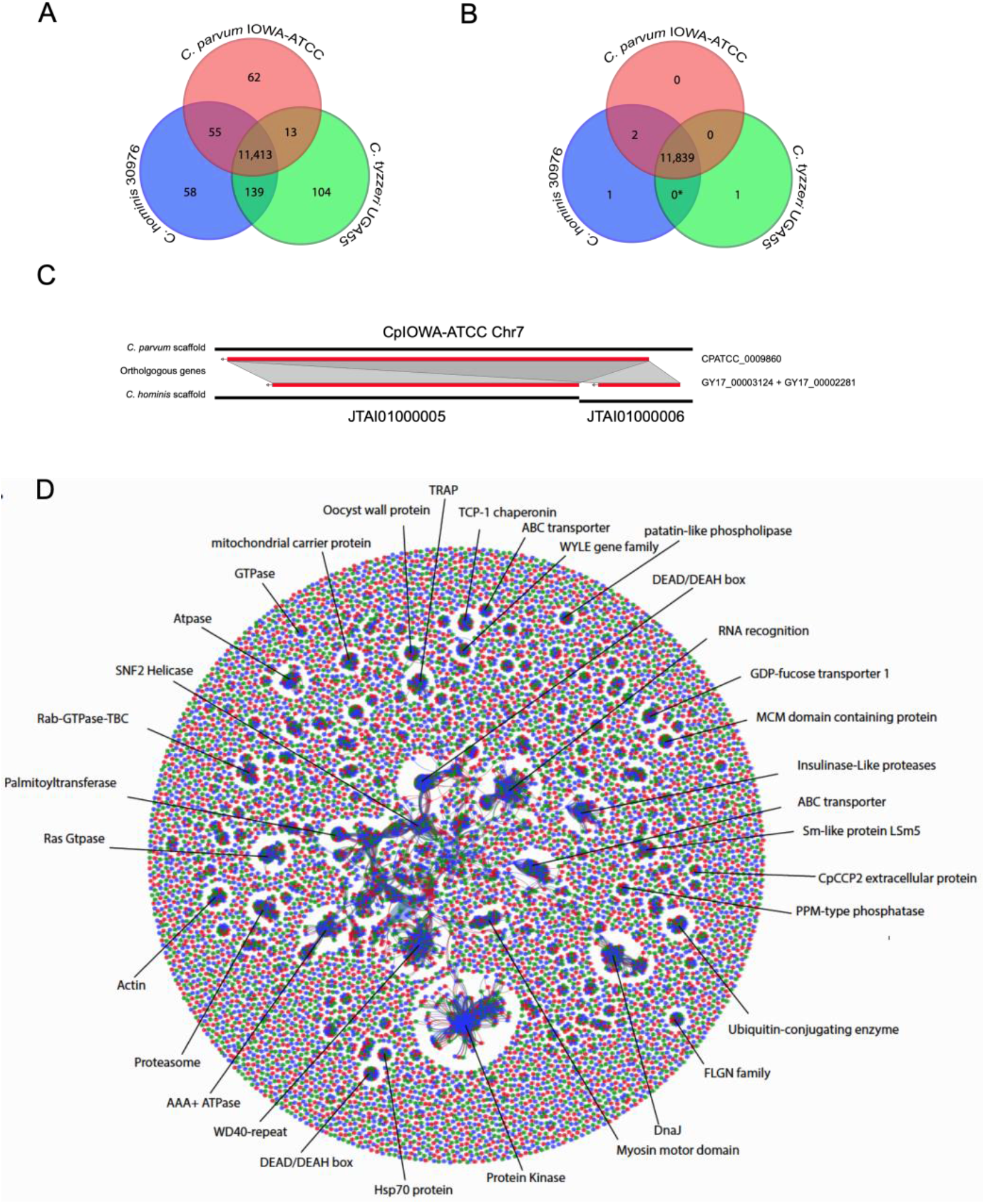
Ortholog distribution of protein-encoding genes. (A) Venn diagram of orthologous gene sequences between three closely related *Cryptosporidium* species (pre-investigation); (B) Venn diagram of same orthologous gene sequences (post-investigation) following removal of false species-specific genes, e.g. artifacts. *The 139 genes shared between *C. hominis* and *C. tyzzeri* in panel A are in complex regions with repeats and gaps and do not have enough evidence to prove their uniqueness at this stage given the available assemblies, so they are considered artifacts at this time; (C) Orthology based synteny overview of Chr7 singleton-like, putative paralog artifact generated by a split gene due to the genome assembly fragmentation in one species; and (D) Graphical representation of ortholog clusters between the three closely related *Cryptosporidium* species. *C. parvum* IOWA-ATCC: red; *C. hominis* 30976: blue; *C. tyzzeri* UGA55: green.

We mapped Illumina reads from *C. parvum* IOWA, *C. hominis* TU502-2012 and *C. tyzzeri* UGA55 to the new *C. parvum* IOWA-ATCC long-read assembly to identify and assess putatively overly collapsed regions (repetitive regions represented by only a single repeat in the assembly) (Supplemental Table S1; Supplemental Fig. S4). Our pipeline detected 14 compressions > 100 bp in length in the *C. parvum* IOWA II genome assembly compared to 8 in the new *C. parvum* IOWA-ATCC assembly. These compressions are not always related to genic regions and vary in genome location and predicted copy number. Some of these apparently collapsed regions, were conserved between both *C. parvum* assemblies but varied in different species (Supplemental Fig. S5). The collapsed genic regions are composed of rRNA genes, some uncharacterized proteins, GMP synthase, aspartate-ammonia ligase, tryptophan synthase beta and MEDLE genes. Most of the observed and fixed compressions do not contain any annotated genes.

### A closer look at subtelomeric regions reveals their complexity and relevant biology

As shown in the read depth coverage analysis and in Supplemental Table S1, the new assembly was able to fix most of the collapsed regions in the *C. parvum* IOWA-ATCC genome. Interestingly, one subtelomeric region in Chr1 still has compressions suggesting that most of the genes present in this region have more than one copy (Fig. 5). This region reveals at least 13 genes which vary in copy number between different *Cryptosporidium* species (Supplemental Fig. S5). The genes contained in this region are 18S rRNA, 5S rRNA and 28S rRNA, uncharacterized proteins, a GMP synthase, an aspartate-ammonia ligase, tryptophan synthase beta and a cluster of several MEDLE genes. Some of these genes, such as the tryptophan synthase beta and the MEDLE’s are the focus of considerable research since they may be related to parasite survival and are potentially involved in parasite invasion, respectively (Sateriale and Striepen 2016; Li et al. 2017; Fei et al. 2018). The number of copies predicted here for the rRNAs and MEDLE’s are underrepresented as they also have paralogs on Chr 2 and Chr 5, respectively.

**Figure 5.**
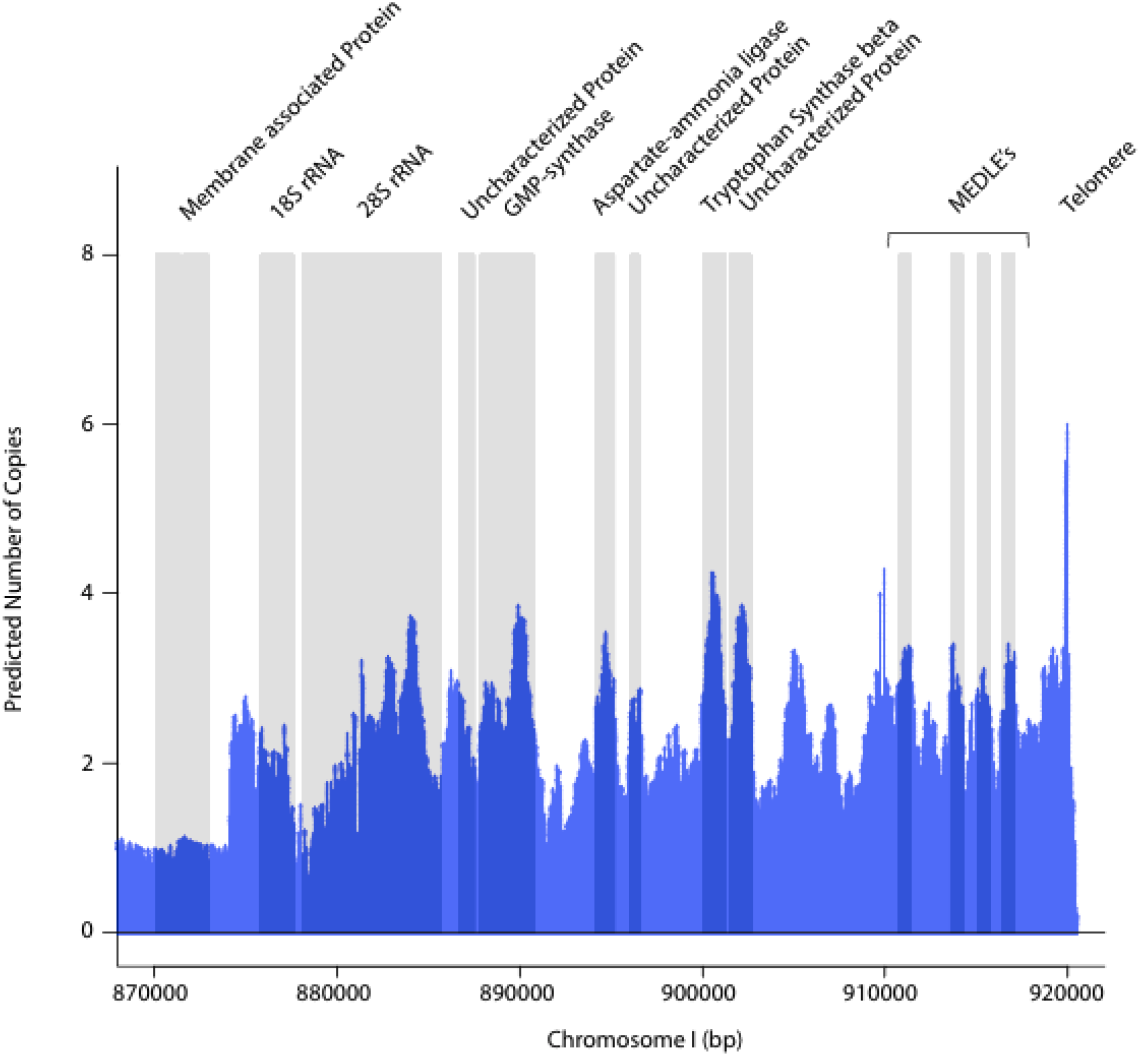
Chromosome 1 subtelomeric region read depth coverage plot normalized by single copy genes. Illumina reads from *C. parvum* IOWA-ATCC DNA are mapped to the *C. parvum* IOWA-ATCC long-read assembly to identify read pileups and estimates of sequence copy number. Vertical grey areas indicate regions with annotated genes. The GMP-synthase shaded region also contains a small uncharacterized protein.

Since we have an apparent compression in a subtelomeric region assembly with no gaps and good PacBio long read coverage, we hypothesized that these extra copies might derive from unassembled regions. The chromosomal-level IOWA-ATCC assembly was only missing three telomeric regions, both ends of Chr 7 and one telomere of Chr 8. Using existing PacBio long-reads we were able to identify a few reads that extended into rRNA regions on the chromosomes missing telomeres. We attempted re-assembly with only PacBio reads and we could not convincingly resolve the missing regions. Thus, we generated very deep (1200 X) Oxford Nanopore (ONT) single molecule reads from *C. parvum* IOWA-BGF (ATCC was not available). The ONT reads revealed related, yet unique subtelomeric regions linked to the chromosomes missing their telomeres, in addition to Chr 1 (Fig. 6). We found good ONT long-read support for these regions. Notably, each different subtelomeric region is flanked by ribosomal RNAs and we also note that there is slight variation observed among the reads for each chromosome end.

**Figure 6.**
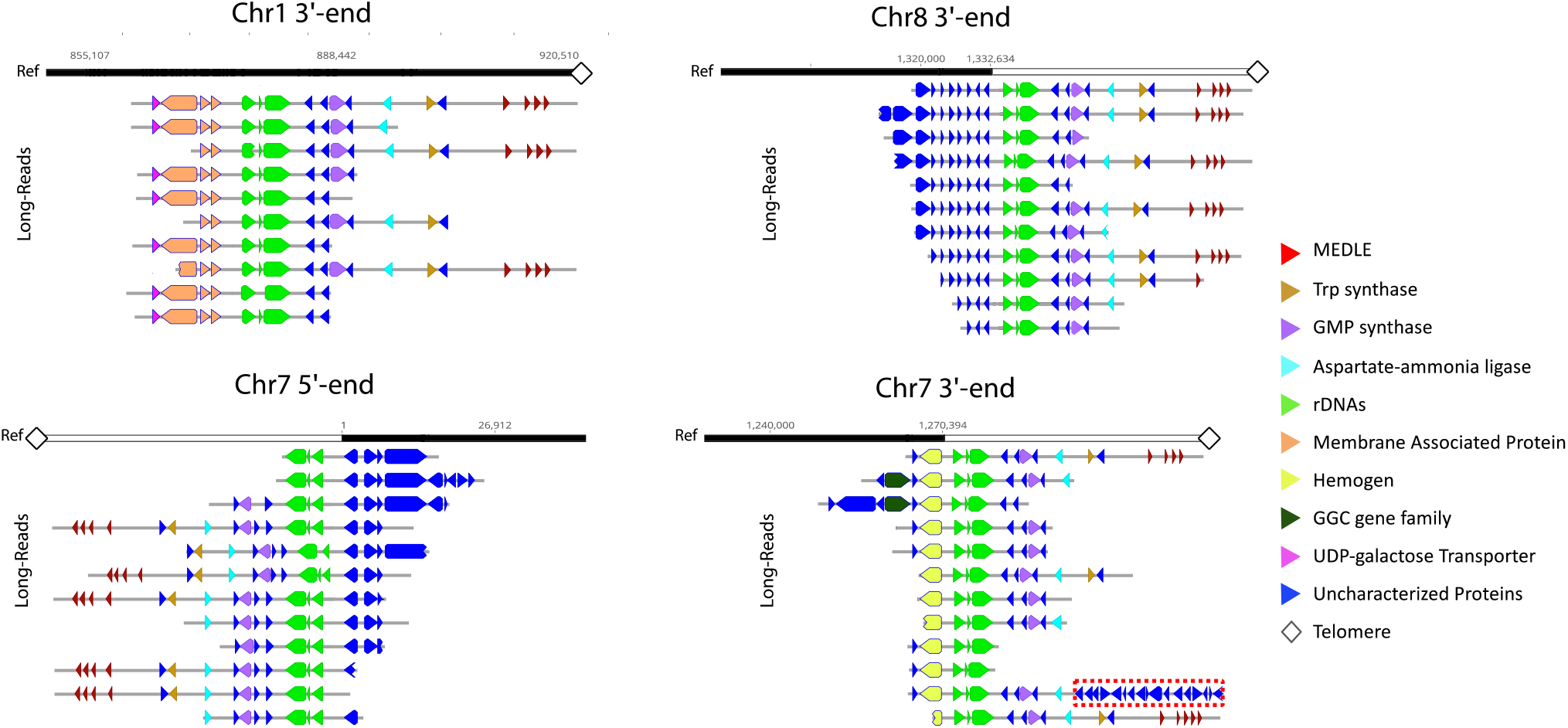
Related subtelomeric regions on different *C. parvum* chromosomes are supported by ONT long-reads. Individual ONT long-reads provide evidence of at least four different, yet related, subtelomeric regions that extend into the chromosomes that were missing telomeres (Chr 7 and Chr 8) in addition to Chr 1. The white and black reference bar above each collection of annotated Nanopore reads identify the newly identified subtelomeric regions (white) and existing assembly (black). The red box on the penultimate read on the Chr 7 3’ end panel indicates a unique region of insertion (nucleotide positions 1191705-1217462). This region contains mostly uncharacterized proteins and two transferases. Each ONT read is annotated as indicated in the key.

### Fast evolving genes in *C. parvum*

The new gapless genome assembly and annotation presented an opportunity to revisit the prediction of fast-evolving genes in this species. We performed a Single Nucleotide Variant, SNV, analysis using 136 different *C. parvum* WGS data sets obtained from GenBank (Supplemental Table S9) using the new assembly and annotation. A total of 24,407 positions were found to contain at least one high-confidence bi-allelic variant. Multiallelic calls were removed to guard against mixed infections. The biallelic variants reflect 3892 genes, 190 of which show a Ka/Ks ratio of non-synonymous/synonymous rates of > 1.0 (Supplemental Table S10). Of the 190, 24 genes were previously identified and 124 are classified as uncharacterized proteins, 93 of which are annotated as having a signal peptide or being secreted. All previously identified fast evolving genes were detected, including: Insulinase-like protein (CPATCC_0017080), an uncharacterized secreted protein (CPATCC_0010380), *gp60* (CPATCC_0012540) and others (Strong et al. 2000; Sanderson et al. 2008; Nader et al. 2019; Zhang et al. 2019). The top eight genes by Ka/Ks ration have not been previously reported. Gene family members such as MEDLEs, FLGN and SKSR were also detected but significantly, new members of each of these families are identified as also under positive selection. A family of WYLE (Sanderson et al. 2008) proteins is also identified as under selection.

## DISCUSSION

The first genome assembly sequence of *Cryptosporidium parvum* IOWA II (Abrahamsen et al. 2004) was excellent given the technology at the time and because of its quality the community has relied on this genome assembly and annotation to design their experiments. However, gaps and ambiguous bases remained, and there was little available expression or orthology evidence to assist the annotation. We used PacBio, Nanopore and Illumina sequencing technologies to generate a new complete genome assembly of *C. parvum* strain IOWA-ATCC. We then applied de novo and evidence-based annotation approaches with manual curation of two additional species to generate consistent annotation that could be used to detect unique genes and genomic differences between species and strains.

The first, expected, finding was that the *C. parvum* IOWA stain is continuing to evolve (Cama et al. 2006) as it is maintained by passage through cattle in a few different locations for research use. Some natural *Cryptosporidium* isolates have been propagated in unnatural hosts before sequencing. Thus, selection during propagation or maintenance via animal propagation may lead differences relative to circulating parasites. This phenomenon has been observed in other protozoan parasites (Lecomte et al. 1992; Sutherland et al. 1996; Akiyoshi et al. 2002; Chan et al. 2015; Isaza et al. 2015). Genomic DNA for the 2004 *C. parvum* IOWA II and *C. parvum* IOWA-ATCC were obtained from the same source, but many years apart. We note small differences in the *gp60* sequence, and an overall genome average difference of ~0.07% in identity (Supplemental Table S2). Changes were also observed in *Plasmodium* species, after being propagated for a long time period (Claessens et al. 2017) and is associated with loss of infectivity and virulence in some strains (Segovia et al. 1992).

When compared to *C. hominis* and *C. tyzzeri*, which are 95–97% identical in available nucleotide sequence, incongruences in the annotated gene models with respect to the new *C. parvum* IOWA-ATCC genome assembly were obvious. The differences result in part from some genome assemblies that contain numerous sequence gaps and little experimental evidence (i.e. RNA-Seq data for all major developmental stages) to permit accurate annotation. The gaps can lead to pseudo-gene annotations or split genes due to frame-shift artifacts. For example, regions with gaps or errors in the base call can lead to false stop codons, or frameshifts that are usually detected as incomplete pseudogenes or a gap can cause a predicted gene to be split into more than one piece. Additionally, *in silico* prediction tools are usually not trained for non-model organisms and *C. parvum* is so distant from other sequenced organisms, there is little synteny or orthology to help guide the various efforts. These mis-annotations can be detected and avoided if an evidence and homology-based curation between different samples is conducted.

As observed in Table 2, annotated protein-coding gene numbers do not exactly match between the closely related species with high sequence identity. This difference is explained by the gaps in the comparator genome assemblies for *C. hominis* and *C. tyzzeri*. These gaps interrupt the open reading frames (ORFs) causing split genes and frame shifts. Thus, some gene models are not necessary missing in an organism. These differences affect similarity-based analyses such as ortholog detection, giving the wrong impression that some of these partially annotated genes are unique for a species (Fig. 4; Supplemental Table S8). These mis-interpretations can sabotage some experimental designs that may use an incorrect basis for experimental design or analysis (Baptista and Kissinger 2019). These regions with problems are usually complex and some have high polymorphism rates (e.g., positive selection). So, false assumptions regarding species-specific genes can affect many downstream analyses including the detection of highly polymorphic loci.

In this study we were able to improve the structural and functional annotation of these genome assemblies, by using two different approaches: (i) using seven full-length stranded cDNA libraries derived from three time points (0h, 24h and 48h post infection) increasing the expression percentage representation of *C. parvum* and transcriptome data from RNA-seq analyses were generated to improve the gene models deposited in CryptoDB.org (Tandel et al. 2019); and (ii) by using homology information to construct a consistent genome annotation between three different close-related species. This approach facilitated a proper comparative analysis of genome content differences between the species compared. Our analyses reveal that the compared species only differ slightly in gene content for the regions that can be compared. Most differences are related to slight structural variation, such as small translocations and inversions, and by copy number variation as revealed by read depth coverage analysis. Previous studies have reported a lack of DNA methylation in *Cryptosporidium* and other parasites (Gissot et al. 2008). The *C. parvum* C-5 cytosine-specific DNA methylase (*Dnmt2*) sequence was previously annotated as truncated (Abrahamsen et al. 2004; Isaza et al. 2015) and lacking a DNMT-specific motif containing a prolyl-cysteinyl dipeptide (Abrahamsen et al. 2004; Ponts et al. 2013; Isaza et al. 2015). The new *Cryptosporidium parvum* IOWA-ATCC whole genome assembly and annotation reveals a complete ortholog of the *Dnmt2* DNA methylase family. The lack of this N-terminus has been cited as a possible reason for the lack of DNA methylation in *C. parvum* (Ponts et al. 2013).

Apicomplexans have reductive streamlined genomes, that range from ~8.5 to ~125 megabases. *Cryptosporidium* species have among the most compacted genomes, with 504 bp average length between the stop codon of one gene and the start codon of the next gene. *Cryptosporidium* also has few protein-encoding genes (~3950) relative to other apicomplexans with up to ~8000 (Kissinger and DeBarry 2011). Studies shows that *Cryptosporidium* may have adapted a novel type of nucleotide transporter for ATP uptake from the host (Pawlowic et al. 2019). Given the compactness of this parasite genome sequence, the gene loss may be compensated for by the higher number of transporters found in our re-analysis. These findings will facilitate future studies of alternative metabolic pathways to better understand the biology and evolution of parasitism of this organism.

Chromosomal inversions are known to affect rates of adaptation, speciation, and the evolution of chromosomes (Guo et al. 2015). Comparative genomic studies and population models for several organisms, suggests that inversions can spread by suppressing recombination between loci and generating areas of linkage disequilibrium. Local adaptation mechanisms applied to demographic and genetic situations, can drive inversion to high frequency if there is no countervailing force, thus explaining fixed differences observed between populations and species (Kirkpatrick and Barton 2006). Previous studies identified potential chromosomal inversion sites between *Cryptosporidium* species relative to *C. parvum* IOWA II (Guo et al. 2015; Isaza et al. 2015). The new long-read genome assembly of *C. parvum* IOWA-ATCC revealed some potential inversion sites, in chr 2, chr 4 and chr 5, that are flanked by poorly sequenced and gapped regions in some species, (Piper et al. 1998; Bankier et al. 2003). Since the other species still lack physical evidence for their chromosomal structures, further long-read sequencing or chromosome conformation capture sequencing, such as Hi-C, is still needed to detect and validate species-specific structural variations for the other *Cryptosporidium* species.

The lack of three telomeres in the new high-quality long-read assembly was an intriguing result that can be explained by the detection of three putative similar but not identical copies of subtelomeric regions containing genes including tryptophan synthase beta, the MEDLE genes and 18S/28SrRNA cluster among others. This finding raises the possibility of this species having misincorporation of telomers by its telomerase, as was observed in other protists (McCormick-Graham et al. 1997) or recombination between telomeres by break-induced replication, such as has been observed in yeasts (McEachern and Iyer 2001; McEachern and Haber 2006), and telomere maintenance by recombination as is observed in human cancers (Natarajan et al. 2006). Since some of the genes in this region are possibly essential genes for parasite survival (Sateriale and Striepen 2016), the fact that they may exists in multiple copies and can possibly generate variation as a result of recombination could explain an alternate new survival mechanism in this streamlined parasite genome. We have support from single molecule sequencing that indeed this region is detected on 4 different chromosome ends (Fig. 6). This potential subtelomeric plasticity resulting in a possible transfer of important gene sequences between homologous and nonhomologous chromosome ends, could affect genetic manipulations and may affect phenotype. We believe that these structures are varying within the *Cryptosporidium* population, which is hard to detect, since we do not yet have any evidence that all 4 related chromosome ends are present in a single cell. Thus, the Nanopore reads may be representing population level variation, which also raises the possibility of recombination or gene conversion as *Cryptosporidium* requires sexual recombination to from excreted oocysts. Currently, cloning does not exist for *Cryptosporidium*. Thus, oocysts used for sequencing must be considered a population even if sequence is derived from single cell sequencing (Troell et al. 2016) as oocysts still contain four haploid meiotic progeny (sporozoites). A truly singe-cell approach, which will facilitate recombination and sub-telomeric plasticity studies, will require single-sporozoite sequencing, but this is still impossible in the absence of genome amplification.

*Cryptosporidium* species are usually typed and characterized by the community using a small number of genetic markers including 18S, COWP, HSP70, and *gp60* (Ghaffari et al. 2014). As shown in this study *gp60* which is a fast-evolving gene used for *Cryptosporidium* subtyping characterization, had small differences between *C. parvum* IOWA II and *C. parvum* IOWA-ATCC. The parasites used to generate these sequences originated from the same propagated strain but were collected at different times. Using just one marker to characterize an obligately sexual organism with 8 chromosomes is problematic. In this study, we confirm an existing group of fast evolving genes and identify 166 additional potential candidates distributed across all 8 chromosomes. Some of these genes belong to gene families so to avoid artifacts only uniquely mapped reads were used for the SNV analysis. The genes identified here can be used to help the community develop additional markers with better resolution for typing parasite isolates. Given that only 136 isolates from a small geographic region have been sampled, the potential to identify additional genes is high. Newer techniques such as hybrid capture bait set techniques (Mamanova et al. 2010) are a powerful future alternative to characterize and select *Cryptosporidium* population variants and better characterize genetic diversity.

The new *C. parvum* long-read assembly combined with a consistent comparative annotation has proven incredibly powerful. The species analyzed here have different host preferences and pathogenicity. Comparisons of previous sequences and annotation suggested numerous gene content differences. However, this systematic study reveals that the primary differences between the zoonotic *C. parvum*, the anthroponotic *C. hominis* and the rodent-infecting *C. tyzzeri* are SNVs and CNVs rather than differences in unique gene content. Finally, new findings related to within parasite and/or within population subtelomeric amplification and variation events in *C. parvum* reveal a new level of genome plasticity that will impact some genetic manipulations and may affect the organisms’ phenotype.

## METHODS

### Sample DNA source and dataset used

*Cryptosporidium parvum* IOWA-ATCC DNA from oocysts/sporozoites was purchased from the ATCC. The source was the University of Arizona, Sterling Parasitology Laboratory. Its GP60 subtype (IIa) is the same as the current *C. parvum* IOWA II reference genome sequence also used in this work. *Cryptosporidium parvum* DNA was also prepared from oocysts obtained in 2018 from Bunch Grass Farms, Deary, ID. This isolate is referred to as IOWA-BGF in this study. The *C. hominis* 30976 and UdeA01 genome assemblies, are human isolates. The *C. tyzzeri* assembly a natural mouse model of Cryptosporidiosis. The 136 *C. parvum* sample accession numbers used for the positive selection analysis are available in Supplemental Table S9.

### *Cryptosporidium parvum* IOWA-ATCC sequencing and genome assembly

PacBio RSII and Illumina HiSeq 2000 sequencing were both performed at the Wellcome Sanger Institute, UK. The *Cryptosporidium parvum* IOWA-ATCC reads were first assembled using the PacBio open source SMRTlink v6.0 from 9 PacBio SMRT cells, with ~75x mean genome coverage. The resulting assembly was then submitted to the accuracy improver tool Sprai 0.9.9.23 (https://sprai-doc.readthedocs.io/en/latest/index.html) and then had gaps filled using PBJelly 15.24.8 (English et al. 2014) using PacBio reads and IMAGE 2.4.1 (Swain et al. 2012) with Illumina reads. A manual inspection and improvement using GAP5 (Bonfield and Whitwham 2010) was needed to better access complex regions, and the final scaffolded genome assembly was polished with Illumina reads using iCORN2 0.95 (Otto et al. 2010) and Pilon 1.22 (Walker et al. 2014).

Oxford Nanopore (ONT) single molecule long-read sequencing was performed on DNA from *C. parvum* IOWA-BGF (The ATCC^®^PRA-67DQ™ ran out of stock) following the protocol recommended by for an R9.4.1 flow cell. MinION ONT sequencing was performed at the Georgia Genomics Bioinformatics Core (GGBC) at the University of Georgia, USA, using an R.9.4 flow cell and the rapid sequencing kit (SKT-RAD004). The ONT long-reads generated >1000x coverage of the *Cryptosporidium parvum* genome. This high coverage complemented the PacBio data to confirm and fix several complex regions. The final assembly was submitted with the current reference and genome assemblies of other closely related species to QUAST v.5.02 (Gurevich et al. 2013) to compare and evaluate the quality of the new genome assembly.

### *Cryptosporidium* genome reannotation

Genome annotation was generated with: (a) an ab initio prediction using GeneMark-ES 4.57 (Lomsadze et al. 2005); (b) evidence-trained predictions by SNAP/Maker (Cantarel et al. 2008; Johnson et al. 2008) and (c) Augustus (Stanke and Morgenstern 2005). For training, we used publicly available data from each respective species: RNA-seq (strand and non-strand specific), ESTs, previously predicted proteins and MassSpec proteomics data when available. In parallel we also generated transcriptome assemblies using HISAT2 v.2.1.0 (Kim et al. 2015) and StringTie v.1.3.4 (Pertea et al. 2015), and non-coding RNA predictions were generated for *C. parvum* as described (Li et al. 2020). Manual curation of all genes in the context of existing molecular evidence was performed using a local installation of WebApollo2 (Lee et al. 2013).

As each genome species analyzed has a different number of publicly available data sets, we also used each curated genome annotation in comparison with the others using the Artemis Comparison tool (ACT) 17.0.1 (Carver et al. 2005), allowing us to perform comparative annotation and resolve discrepancies via homology. All protein-encoding genes annotated for each genome sequence were submitted to OrthoFinder v.2.3.7 (Emms and Kelly 2015) to detect paralogs, orthologs and singletons. All singletons were then selected for a comparative manual curation using MCScanX 0.8 (Wang et al. 2012) and JBrowse (Buels et al. 2016) between all three species to verify their uniqueness and assess the contribution of sequence gaps or misassembly to the findings. We considered the following error types: Split genes caused by frameshifts or early stop-codons, lack of stranded RNAseq to confirm the gene model, and the presence of a gapped region in the genome assembly. All genes that did not fall into one of these categories were considered to be unique.

### Functional annotation

Following structural annotation, the predicted protein sequences were used to search Swiss-pro curated (sprot) and not-curated (Trembl) and the NCBI non-redundant Protein database with BLASTP and an e-value threshold at the superfamily level of 1e-6. Protein structure similarity was explored using I-TASSER (Roy et al. 2010). Protein sequences were divided into two major groups distributed according their length for the I-TASSER analysis: (i) peptide sequences < 750 aa; and (ii) shorter sequential segmental peptide sequences < 750 aa derived from annotated proteins > 750aa. Structures were predicted for each peptide using the I-TASSER suite and aligned to solved crystal structures in the protein data bank (PDB) using the cofactor algorithm (Roy et al. 2012). InterPro codes were assigned to the query peptide sequence via InterProScan v.5.23-62.0 (Quevillon et al. 2005) using 11 different default databases. The PFAM codes available for PDB crystal structures were transposed to InterPro codes using the R pfam.db library. The presence of at least one matching InterPro code assigned to both the query and the reference peptides was taken to indicate a greater likelihood of structural similarity of the predicted structure and considered “high-confidence”. A random forest classifier was trained to distinguish between a test set of high- and low-confidence models, and was then applied to the entire predicted proteome to identify additional high-confidence-like models among unannotated proteins, as described in Ansell et. al 2019 (Ansell et al. 2019). BLAST2GO (Conesa et al. 2005) version 4.1.9 was used to assign Enzyme Code (E.C) and Gene Ontology (GO) terms. Following this functional annotation, we compared the existing protein product names to the new functional results. Some structural information, such as protein domain and repeat pattern content were added to some uncharacterized proteins and nomenclature errors were corrected according to the NCBI annotation submission guide.

### Transporter prediction

Predicted proteins were submitted to four different transporter prediction methods: (i) local alignment using BLASTP against TCDB (Saier et al. 2009) transporter proteins with a threshold e-value of 1e-5 cutoff to find potential transporter similarities; (ii) TMHMM (Server v. 2.0) (Krogh et al. 2001) and SignalP (Server 4.1) (Bendtsen et al. 2004) was applied to reduce false positives from the TCDB blast results. Transporter candidates with no transmembrane domains or candidates with only one transmembrane prediction while having signal peptides predicted were removed; (iii) TransAAP (Ren et al. 2007), which is a TC-based (Transporter Classification from TCDB) transporter annotation tool on the TransportDB v2.0 website (Ren et al. 2007), that was used to provide information about potential transporter identity and substrate; and (iv) a structural proof for candidate transporters using Phyre2.0 (Kelley et al. 2015). Final candidate transporters were checked according to above results as well as annotations obtained from InterProScan 5.44 (Jones et al. 2014).

### Comparative and phylogenetic analysis

Comparative genome-wide synteny between *Cryptosporidium* species was performed using Murasaki v.1.68.6 (Popendorf et al. 2010) with default settings. The cladogram topology was determined via a maximum likelihood analysis of 2700 single copy orthologs using JTT+I as the substitution model as predicted by Modeltest-NG (Darriba et al. 2020). The consensus tree was constructed from 1000 bootstrap replicates. The consistency of annotation and potential gene family copy number variations (CNVs), were determined with Orthofinder v.2.2.7 (Emms and Kelly 2015) which identified all orthologs and paralogs. Orthofinder BLASTP results were parsed to examine the relationships between proteins using an e-value threshold of 1e-20 and identities > 35% between protein pairs longer than 100 amino-acids. The data were visualized using Gephi (https://gephi.org/) with the Fruchterman-Reingold layout.

Copy number variation was also determined by aligning Illumina sequence reads from each closely related species studied to the new *C. parvum* IOWA-ATCC reference genome sequence to check for potential CNV regions by looking for variations in read depth coverage. The alignment was performed using BWA mem 0.7.17 (Li and Durbin 2009) with default options and the alignment depth per base was calculated using BEDTools genomecov 2.29.2 (Quinlan and Hall 2010) and SAMtools depth 1.6 (Li et al. 2009).

### Resolving the structure of repetitive subtelomeric regions

Following the CNV analysis, the sequence content of the putatively compressed regions and their non-compressed sequence boundaries of the *C. parvum* IOWA-ATCC assembly were used to build a BLAST database. We then selected single oxford nanopore single molecule reads using BLASTn 2.10.0 (Camacho et al. 2009) to detect sequences capable of aligning to compressed regions and then determine their putative assembly structures. Following ONT read selection, the ONT reads were polished with Illumina reads using proovread 2.14.1 (Hackl et al. 2014) and Pilon 1.22 (Walker et al. 2014). To map these polished reads against the genome assembly and avoid bias/competion between sites, all putatively compressed genome assembly regions were artificially split into fragments, effectively making the chromosomes with compressed regions fragmented. Reads were aligned to all chromosome fragments using the Geneious mapper 2019.1.3 (https://www.geneious.com) with medium-sensitivity and those chromosome fragments with hits were annotated and analyzed for validation and verification of their structure.

### Variant analysis, selection prediction and populational analysis

Illumina sequence reads from 136 different isolates of *C. parvum* from different geographical locations (Supplemental Table S9) were aligned against the *C. parvum* IOWA-ATCC reference genome sequence using BWA-MEM (Li and Durbin 2009), the bam files were parsed to select uniquely mapped reads and to mark duplicates and remove redundancy using PICARD (Broad_Institute) and then submitted to a Variant call analysis using GATK 3.8 Haplotypecaller (McKenna et al. 2010). These results were then filtered by mapping quality > 40 and depth coverage >10. Because mixed infections exist, we restricted analysis to biallelic sites. The individual VCF files were combined into one GVCF file using the GATK tool GenotypeGVCF. After selecting just single nucleotide variants (SNVs) from this data, the combined gvcf file was annotated using the software snpEff v.4.3 (Cingolani et al. 2012). The number of synonymous and non-synonymous variants were taken from the annotated gvcf file and parsed to calculate the Ka/Ks ratio of non-synonymous/synonymous rates. Genes with ratios > 1.5, indicative of positive selection, were detected and denoted as fast evolving genes within the *C. parvum* population.

## DATA ACCESS

The sequencing data, genomes and annotation generated in this study have been submitted to the NCBI BioProject database (https://www.ncbi.nlm.nih.gov/bioproject/) under accession numbers PRJNA573722, PRJNA252787, PRJEB3213 and PRJNA388495. *C. hominis* UdeA01 assembly and TU502 Illumina reads used are in BioProjects PRJEB10000 and PRJNA222836, respectively. The data are also available at CryptoDB.org (Heiges et al. 2006).

## ACKNOWLEDGMENTS

This work was supported by Bill and Melinda Gates Foundation grant OPP1151701 to JCK, The Wellcome Trust via its core funding of the Wellcome Sanger Institute (grant WT206194) and NHMRC Investigator Grant (APP1194330) to ARJ.

## AUTHORS CONTRIBUTIONS

RPB and JCK designed research; RPB and JCK performed research; AS, JD and BS contributed with new reagents and samples; BA and AJ contributed with analytical tools; MS, KB, AT, MB and JAC contributed Illumina and PacBio sequencing; RPB, YL, KB, AT, RX, EDS, GWC and JCK analyzed data; RPB and JCK wrote the paper and ARJ, BREA, BS, AS and JAC provided feedback.

