## Supplemental Figures for "Long-read assembly and comparative evidence-based reanalysis of *Cryptosporidium* genome sequences reveal new biological insights"

**Affiliation:** [1]Center for Tropical and Emerging Global Diseases; [2]Institute of Bioinformatics, University of Georgia, Athens, GA, USA; [3]Department of Pathology, School of Veterinary Medicine, University of Pennsylvania, Philadelphia, PA, USA; [4]The Wellcome Sanger Institute, Hinxton, UK; [5]Faculty of Veterinary and Agricultural Sciences, The University of Melbourne and Population Health and Immunity Division, the Walter and Eliza Hall Institute of Medical Research, Melbourne, Australia and [6]Department of Genetics, University of Georgia, Athens, GA, USA

### Supplemental Methods:

#### Multiple sequencing Alignment (Figs. S1, S5 and Table S1)

Multiple sequencing alignment of *Cryptosporidium gp60* and 5C-cytosine-specific DNA methylase was performed by MAFFT v.7 (Katoh and Standley 2013) using default parameters. Sequences were retrieved from CryptoDB v.49, PlasmoDB v.49 and BLAST in NCBI GenBank. Alignments were visualized using plain text Clustal format and Geneious Prime 2019.1.3 (Kearse et al. 2012).

#### Evaluation of compressed regions and CNV genes among the three close related species (Fig. S2, Fig. S4)

For Illumina read depth coverage (RDC) analysis and detection of CNVs among the three different species, *C, parvum* IOWA-ATCC, *C. hominis* TU502 and *C. tyzzeri* UGA55, the alignment was performed using BWA mem 0.7.17 (Li and Durbin 2009) with default options and the alignment depth per base was calculated using BEDTools genomecov 2.29.2 (Quinlan and Hall 2010) and SAMtools depth 1.6 (Li et al. 2009). *C. hominis* was represented by a different strain (TU502_2012) in this analysis since the *C. hominis* 30976 strain Illumina sequencing library was generated by an amplification selection method (WGA), which would affect the read depth coverage analysis. For a better whole genome overview of any putatively compressed regions, with potential CNV we used the data.table R package with a sliding window of 100 bp and only selected regions > 2 predicted collapsed window copies. The coverage plots were generated using Circa (http://omgenomics.com/circa).

All regions with higher coverage were compared to the GFF file using BEDTools for gene detection and had their copy number estimated by dividing their average RDC by the estimated single copy coverage of the sequencing.

**Supplemental Figures:**

```
Cparvum_ATCC     MRLSLIIVLLSVIVSAVFSAPAVPLRGTLKDVPVEGSSSSSSSSSSSSSSSSSSSTSTVA
Cparvum_IOWAII   MRLSLIIVLLSVIVSAVFSAPAVPLRGTLKDVPVEG--SSSSSSSSSSSSSSSSSTSTVA
                 ************************************  **********************

Cparvum_ATCC     PANKARTGEDAEGSQDSSGTEASGSQGYEEEGSEDDGQTSAASQPTTPAQSEGATTETIE
Cparvum_IOWAII   PANKARTGEDAEGSQDSSGTEASGSQGSEEEGSEDDGQTSAASQPTTPAQSEGATTETIE
                 *************************** ********************************

Cparvum_ATCC     ATPKEECGTSFVMWFGEGTPAATLKCGAYTIVYAPIKDQTDPAPRYISGEVTSVTFEKSY
Cparvum_IOWAII   ATPKEECGTSFVMWFGEGTPAATLKCGAYTIVYAPIKDQTDPAPRYISGEVTSVTFEKSD
                 ***********************************************************

Cparvum_ATCC     NTVKIKVNGQDFSTLSANSSSPTENGGSAGQASSRSRRSLSEETSEAAATVDLFAFTLDG
Cparvum_IOWAII   NTVKIKVNGQDFSTLSANSSSPTENGGSAGQASSRSRRSLSEETSEAAATVDLFAFTLDG
                 ************************************************************

Cparvum_ATCC     GKRIEVAVPNVEDASKRDKYSLVADDKPFYTGANSGTTNGVYRLNENGDLVDKDNTVLLK
Cparvum_IOWAII   GKRIEVAVPNVEDASKRDKYSLVADDKPFYTGANSGTTNGVYRLNENGDLVDKDNTVLLK
                 ************************************************************

Cparvum_ATCC     DAGSSAFGLRYIVPSVFAIFAALFVL
Cparvum_IOWAII   DAGSSAFGLRYIVPSVFAIFAALFVL
                 **************************
```

**Supplemental Figure S1.** GP60 protein sequence alignment between *C. parvum* IOWA-ATCC and *C. parvum* IOWA II. The amino acid differences are highlighted.

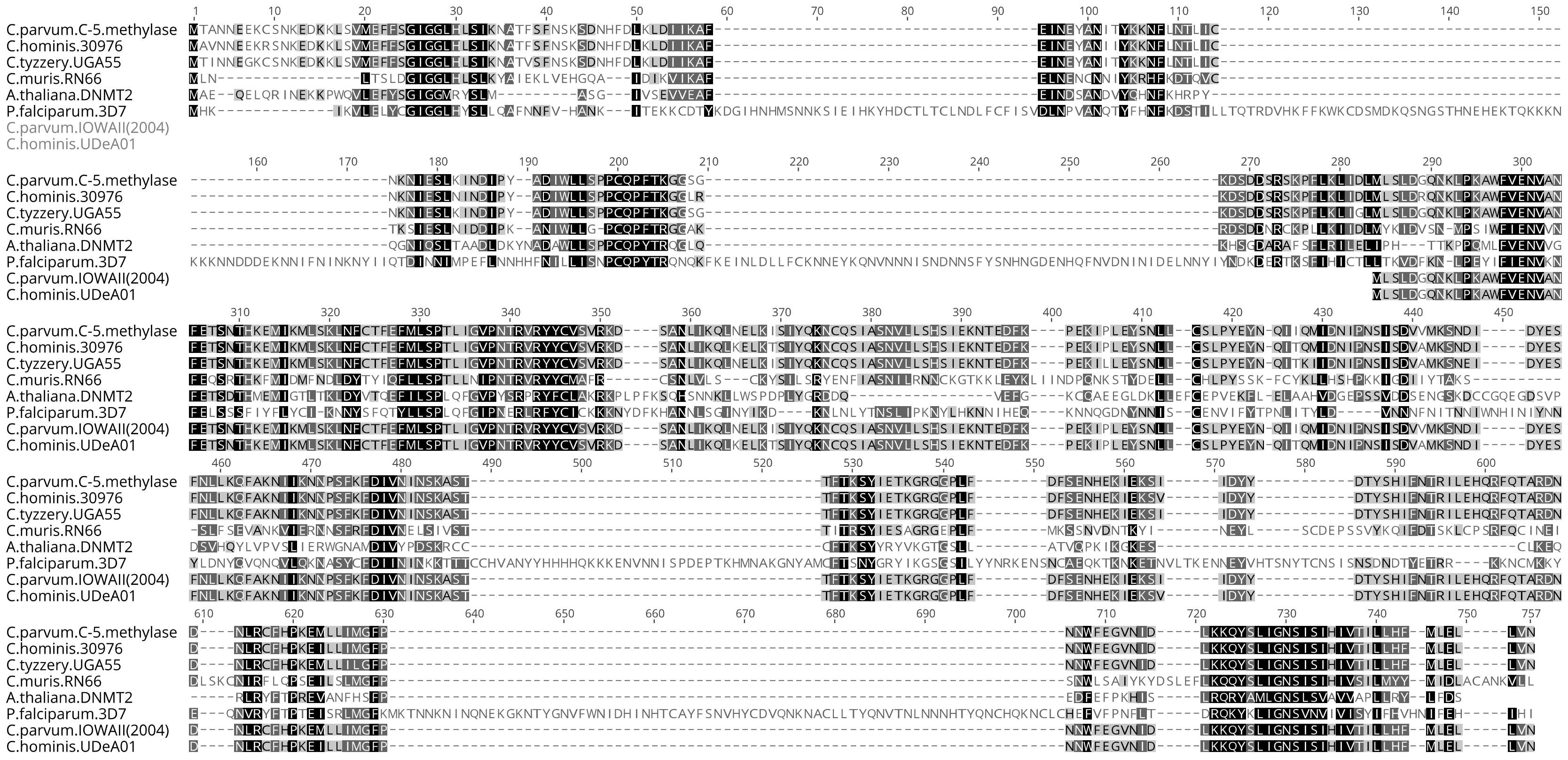


**Supplemental Figure S2.** C-5 cytosine-specific DNA methylase alignment between the reannotated *Cryptosporidium* species compared to some other species and the older truncated annotation available.

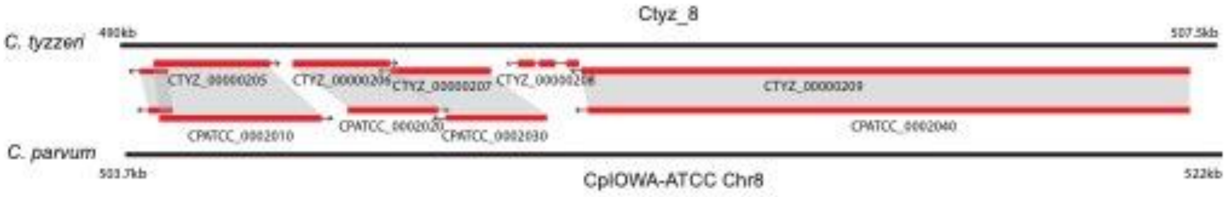


**Supplemental Figure S3.** Orthology based synteny overview of the *C. tyzzeri* potential species-specific uncharacterized protein found on Chr8 (CTYZ_00000208). Grey shading indicates orthology between *C. tyzzeri* and *C. parvum* genes shown in red. The chromosomes, shown in black do not contain any gaps in this region. Figure reproduced based on JBrowse genome viewer.

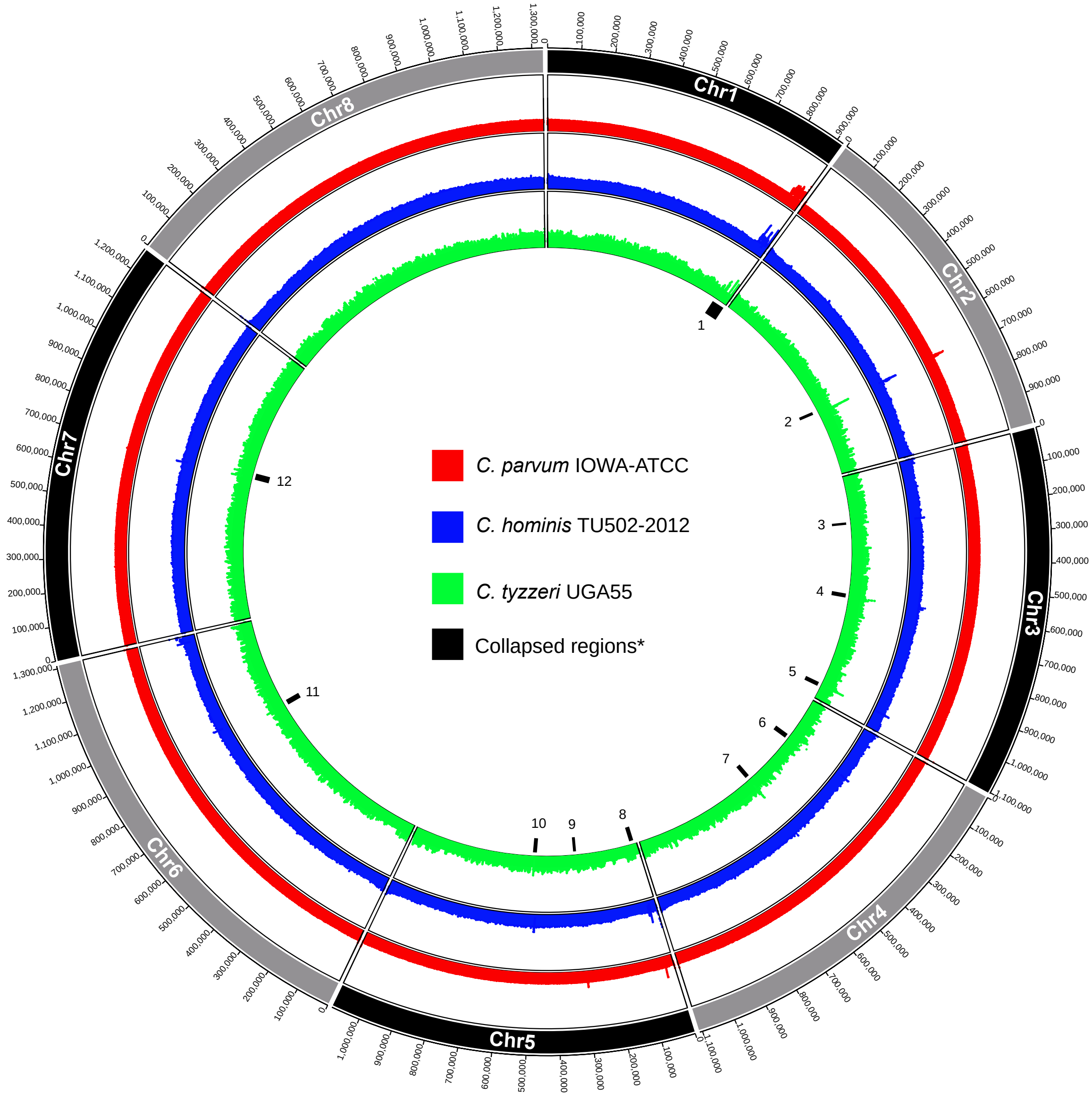


**Supplemental Figure S4.** Read depth coverage distribution among the three closely related species of *Cryptosporidium* using *C. parvum* IOWA-ATCC as reference. The coverage was made using a 100 bp sliding window. *The 12 regions (black tick marks) annotated with numbers on the innermost circle identify putatively collapsed regions with > 2X the level of overall genome read coverage. Numbers 1 and 2 correspond the regions with annotated genes described in the text.

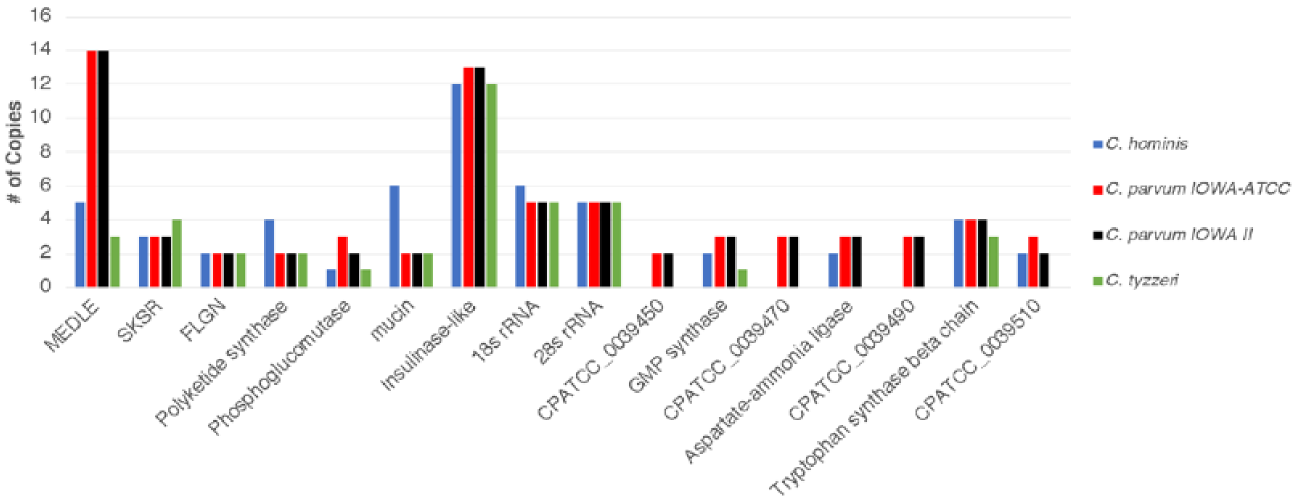


**Supplemental Figure S5**. Genes located in putatively compressed regions of the new *C. parvum* IOWA-ATCC assembly. The major gene families are highlighted. Genes IDs are used to represent uncharacterized annotated protein families.
